# Inter-areal balanced amplification enhances signal propagation in a large-scale circuit model of the primate cortex

**DOI:** 10.1101/186007

**Authors:** Madhura R. Joglekar, Jorge F. Mejias, Guangyu Robert Yang, Xiao-Jing Wang

## Abstract

Reliable signal transmission represents a fundamental challenge for cortical systems, which display a wide range of weights of feedforward and feedback connections among heterogeneous areas. We re-examine the question of signal transmission across the cortex in network models based on recently available mesoscopic, directed‐ and weighted-inter-areal connectivity data of the macaque cortex. Our findings reveal that, in contrast to feed-forward propagation models, the presence of long-range excitatory feedback projections could compromise stable signal propagation. Using population rate models as well as a spiking network model, we find that effective signal propagation can be accomplished by balanced amplification across cortical areas while ensuring dynamical stability. Moreover, the activation of prefrontal cortex in our model requires the input strength to exceed a threshold, in support of the ignition model of conscious processing, demonstrating our model as an anatomically-realistic platform for investigations of the global primate cortex dynamics.

---

In computational neuroscience, there is a dearth of knowledge about multi-regional brain circuits. New questions that are not crucial for understanding local circuits arise when we investigate how a large-scale brain system works. In particular, reliable signal propagation is a prerequisite for information processing in a hierarchically organized cortical system. A number of studies have been devoted to signal propagation from area to area in the mammalian cortex [1–8]. It was found that a major challenge is to ensure stable transmission, with the signal undergoing neither successive attenuation nor amplification as it travels across multiple areas in a hierarchy. In spite of insights these studies provided, virtually all previous works did not incorporate data-constrained cortical connectivity, and made unrealistic assumptions – for instance, areas are considered identical, network architecture is strictly feedforward, and connection weights are the same at all stages of the hierarchy. Here we argue that achieving stable signal propagation becomes even more challenging with the inclusion of more realistic network architecture and connectivity. Mechanisms that improve signal propagation in simpler models may no longer work for more biological models.

Inter-areal cortical network is highly recurrent, abundant with feedback loops [9]. These rich feedback connections pose the risk of destabilizing the system through reverberation as the signal is transmitted across areas. Local cortical circuits are strongly recurrently connected [10], further contributing to system instability. Therefore, mechanisms that improve signal propagation in feedforward networks may quickly lead to instability in a brain-like network. Building such a model requires quantitative anatomical data. Recently, detailed mesoscopic connectivity data has become available for both macaque monkey [9–11] and mouse [12, 13]. This data delineates a complex inter-areal cortical network with connection weights spanning several orders of magnitude. Some areas receive strong inter-areal connections while some other areas appear more disconnected from the rest of the cortex. It is particularly challenging to facilitate signal propagation to the weakly connected areas while maintaining stability for the more strongly connected hub areas [14]. Recent work employs directed‐and weighted inter-areal connectivity data to build biologically realistic large-scale circuit dynamical models of the primate cortex [15, 16], with both local and long-range recurrent connections. This connectivity data spans 29 widely distributed cortical areas, across the occipital, temporal, parietal, and frontal lobes [9]. These anatomically calibrated models thus provide a useful framework for re-examination of signal propagation in the cortex.

We propose a novel biologically plausible mechanism to improve reliable cortical signal transmission. Our mechanism is inspired by the balanced amplification mechanism [17], extending its central idea from the local circuit to a large-scale system. Balanced amplification follows from an underlying circuit connectivity characterized by strong recurrent excitation stabilized by inhibition. We test our mechanism in a range of large-scale models of the primate cortex, including recent population-rate models with heterogeneity across areas [15] and with a cortical laminar structure [16]. In order to examine synchronous [5–7] and asynchronous [3, 4, 18, 19] transmission, which cannot be properly captured with firing rate models, we build a large-scale cortical spiking network model, the first of its kind. We find that our mechanism improves signal transmission by up to 100 fold in all these models. Furthermore, our large-scale spiking model displays several key features of subliminal, preconscious, and conscious processing [20, 21]. Taken together, the findings demonstrate that our network models offer a valuable platform to study a wide range of dynamical questions that involve long-range interactions between cortical areas.

## Results

### Transmission in a realistic large-scale cortical network

Here we demonstrate the challenge of reliable signal transmission through a large-scale network constituted by population rate models [15], where the inter-areal connectivity is set according to a connectivity dataset of the macaque cortex [9]. The directed‐and weighted connectivity matrix was obtained using tract-tracing techniques [9] (see Methods). Briefly, a retrograde tracer was injected into a given (target) area, labeling presynaptic neurons in source areas that connect to the target area. The relative weight of a directed connection was measured as the number of labeled neurons in a source area divided by the total number of labeled neurons in all source areas, called Fraction of Labeled Neurons (FLN) [9]. Heterogeneity was introduced across cortical areas [15], assuming the number of spines per pyramidal cell as a proxy of the strength of excitatory inputs that varies from area to area [22].

On investigating signal transmission in a dynamical model of large-scale macaque cortex [15], we find that inter-areal excitatory loops between cortical areas make reliable signal transmission especially difficult. In Fig. 1A, an input is applied to V1, which is the lowest in the cortical hierarchy [15, 23] (see Methods), and the maximum firing rate is shown for V1 and area 24c at the top of the hierarchy. Inter-areal connections in the model are governed by two global coupling parameters *μ*_*EE*_ and *μ*_*IE*_, corresponding to the long-range excitatory-to-excitatory and excitatory-to-inhibitory coupling, respectively. A small increase in *μ*_*EE*_ can result in the system behavior switching from strong attenuation to instability, as shown in Fig. 1B. A more systematic characterization of the model’s behavior reveals that a gradual increase in *μ*_*EE*_ leads to a sharp transition from a regime characterized by strong signal attenuation to a regime indicating instability (Fig. 1C). Neither regime allows for a realistic propagation of the signal across cortical areas.

**Fig. 1:**
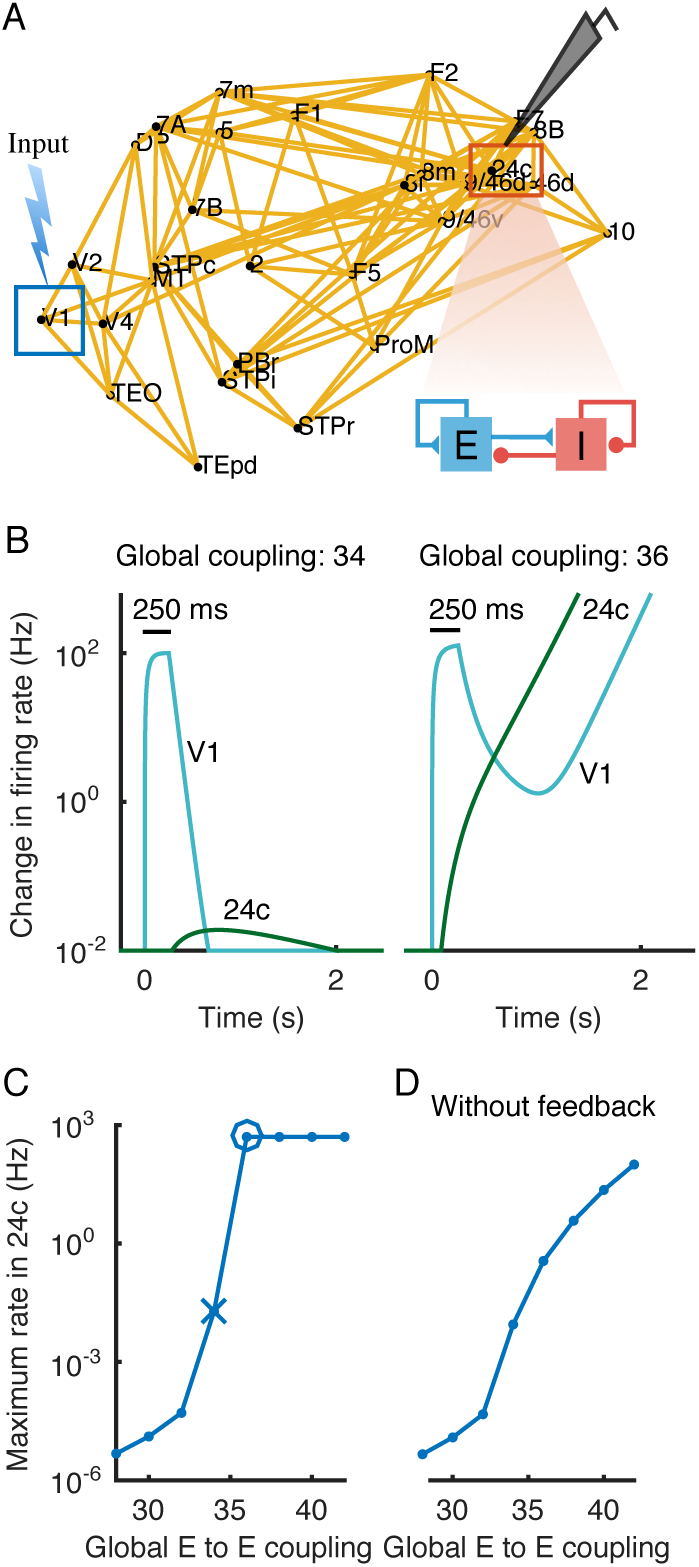
Signal propagation in a large-scale network with recurrent excitatory loops leads to either strong attenuation or instability. (A) Network model of the 29 cortical areas with the strongest inter-areal projection strengths (FLN values > 0.005). Input is injected to the excitatory population in area V1 at the lowest position in the cortical hierarchy, and the peak firing rate in area 24c at the top of the hierarchy is recorded. (B) The response of the excitatory populations in areas V1 and 24c to a current pulse of 250 ms to V1, using two close values of the excitatory global coupling parameter *μ*_*EE*_. (Left) Response in 24c shows an attenuation of 4 orders of magnitude when *μ*_*EE*_ = 34. (Right) With *μ*_*EE*_ = 36, excitatory firing rates in V1 and 24c exponentially grow, and the system becomes unstable. (C) The peak excitatory population response in 24c as a function of *μ*_*EE*_, dynamical instability corresponds to capped firing rate at 500 Hz. The particular parameter values corresponding to weak propagation and instability in (B) are indicated by a cross and a circle respectively. (D) Similar to (C), but *μ*_*EE*_ is varied in the absence of feedback projections.

To answer the question of whether this sharp transition is due to the inter-areal excitatory loops, we examine the model’s behavior when feedback projections are removed from the network. This reveals a smooth transition from the regime with weak propagation to a regime with improved propagation (Fig. 1D), suggesting that removing feedback projections, and therefore the presence of inter-areal excitatory loops, alleviates the problem of effective transmission. Around half of the inter-areal projections present in the anatomical connectivity data, however, correspond to feedback projections, with strengths comparable to those of feedforward projections [9] (Fig. S1), which suggests that feedback projections cannot be ignored. The question of signal propagation, therefore, becomes especially pertinent in a biologically realistic cortical model, endowed with feedback connections.

### Extending balanced amplification beyond the local circuit

To enhance signal propagation while maintaining system stability, we propose a mechanism of transient signal amplification. The mechanism for transient amplification was originally studied in local inhibition-stabilized network models [17, 24]. These networks are characterized by a strong recurrent excitation, which drives the neural activity towards instability, followed by a strong lateral inhibition which stabilizes neural activity (Fig. 2A). These two factors combined, result in a transient amplification of the excitatory firing rate in response to a brief input prior to stabilization, a phenomenon referred to as balanced amplification [17] (Fig. 2B), or Local Balanced Amplification (LBA) for a local network. Increasing LBA can evoke a stronger transient excitatory response prior to decay (Figs. 2B, C), which leads to a transient amplification of activity in the local circuit [17]. It can be analytically shown, using a phase diagram of the network activity as a function of connectivity parameters, that moving along the stability boundary in the direction of increasing LBA (Fig. 2C) leads to a progressive increase in the steady-state excitatory firing rate (Supplementary Appendix). This can be used to intuitively understand the transient amplification achieved with stronger LBA.

**Fig. 2:**
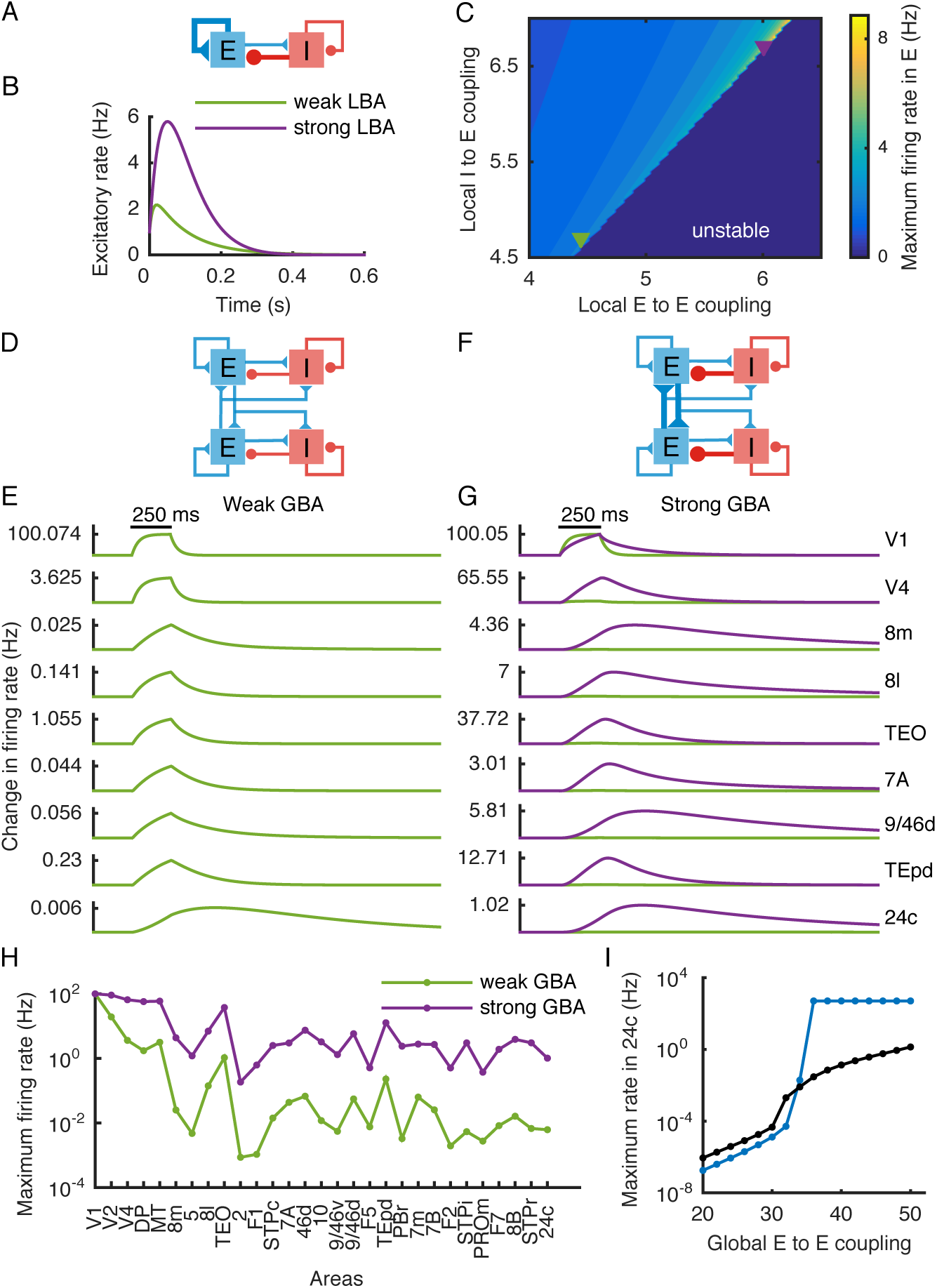
Global Balanced Amplification in the large-scale model improves signal propagation. (A) Model scheme with strong recurrent excitation balanced by strong lateral inhibition. (B) Local Balanced Amplification (LBA) results in a transient amplification of the excitatory firing rate prior to decay, in response to a brief input that sets the initial rate to 1. (C) Peak response of the excitatory firing rate as a function of recurrent excitation (local Eto-E connection) and lateral inhibition (local I-to-E connection). The blue region on the upper right indicates instability. The green and purple triangles correspond to the parameter values used in (B). (D-E) With weak GBA there is 10000-fold attenuation of signals from V1 to area 24c (top to bottom). (F-G) With strong GBA, signal propagation is enhanced by 100 fold (purple), overlaid on the green curve from (E). (H) The maximum firing rate across areas as the response to the pulse input to V1 propagates along the hierarchy, for weak (green) and strong (purple) GBA. (I) Peak firing rate response in area 24c with strong GBA (black) overlaid on the curve corresponding to a sole increase in global excitatory coupling (blue curve, Fig. 1C), demonstrating that network instability is prevented by GBA.

A key idea of the present work is an extension of this mechanism from local circuits to large-scale models (Fig. 2D), to boost inter-areal signal transmission. To this end, we replace the increase in local recurrent excitation by an increase in the global excitatory coupling *μ*_*ΕΕ*_, and stabilize the system with stronger lateral inhibition, as in the case of LBA. This principle of strong long-range excitation, stabilized by strong local inhibition, constitutes an extension of the balanced amplification mechanism for large-scale systems, which we term as Global Balanced Amplification (GBA). From now on, we refer to increasing global excitatory coupling and stabilizing the system with stronger lateral inhibition, while keeping other model parameters the same, as increasing GBA.

To quantify how increasing GBA affects propagation in the large-scale network model [15], we measure the quality of signal transmission by comparing the peak value of the excitatory firing rate in area 24c with the same peak value in area V1; the ratio between both peaks is defined as the “propagation ratio”. The response of the different cortical areas to a pulse input in V1, as the signal propagates along the hierarchy, shows a strong attenuation of ~10,000 fold [15] (Fig. 2E). Interestingly, by increasing GBA (Fig. 2F), the propagation ratio is improved by around two orders of magnitude (Fig. 2G). More precisely, a substantial improvement is observed across most of the cortical areas (Fig. 2G,H). More systematic simulations show that, as opposed to simply increasing the global excitatory coupling (Fig. 1C), increasing GBA leads to a smooth transition from the weak to the improved propagation regime (Fig. 2I).

Our mechanism is robust when feedback projections are removed from the large-scale network or in the absence of heterogeneity across areas (Fig. S2A, B). Moreover, we test our mechanism after symmetrizing the anatomical connectivity matrix, and find that the improvement in propagation doesn’t depend on the specific directionality of connectivity (which is not captured by the commonly used diffusion tensor imaging method) (Fig. S2C, D). Inspired by the “small-world” property characterized by high clustering coefficients and short path-lengths, we assess the effect of removing weak anatomical connections with a parametric threshold of connection strength [25] (Fig. S3). In all these cases, the mechanism consistently reveals a significant improvement in propagation with stronger GBA.

From a mathematical point of view, balanced amplification in inhibition-stabilized networks results from the non-normality of the underlying connectivity matrix [17] (a non-normal matrix is one of which eigenvectors are not mutually orthogonal). Non-normality can be examined through Schur decomposition, by expressing the effective connectivity across network basis patterns through self-connections and feedforward connections. An improvement in signal propagation in the model correlates with an increase in the non-normality measure [26] of the anatomical connectivity matrix underlying the large-scale dynamics (Fig. S4).

### Incorporating a cortical laminar structure

Does our signal propagation mechanism still hold in a cortical system where feedforward and feedback projections are wired in a layer dependent manner [23, 27, 28]? We further test efficient signal propagation, using a recent model which incorporates laminar structure in the cortical areas [16] (Fig. 3A). Increasing GBA improves signal transmission in this model as well (Fig. 3B), suggesting that our mechanism is also valid for models equipped with more detailed laminar-specific projections patterns [16]. Improved propagation is accompanied by a small change in gamma power across areas (Fig. 3B), stemming from the stronger lateral inhibition associated with improved GBA. The moderate gamma power is in good agreement with experimental observations, which show the presence of a weakly coherent gamma rhythm during cortical interactions, and its importance in local and long-distance interactions [29–31].

**Fig. 3:**
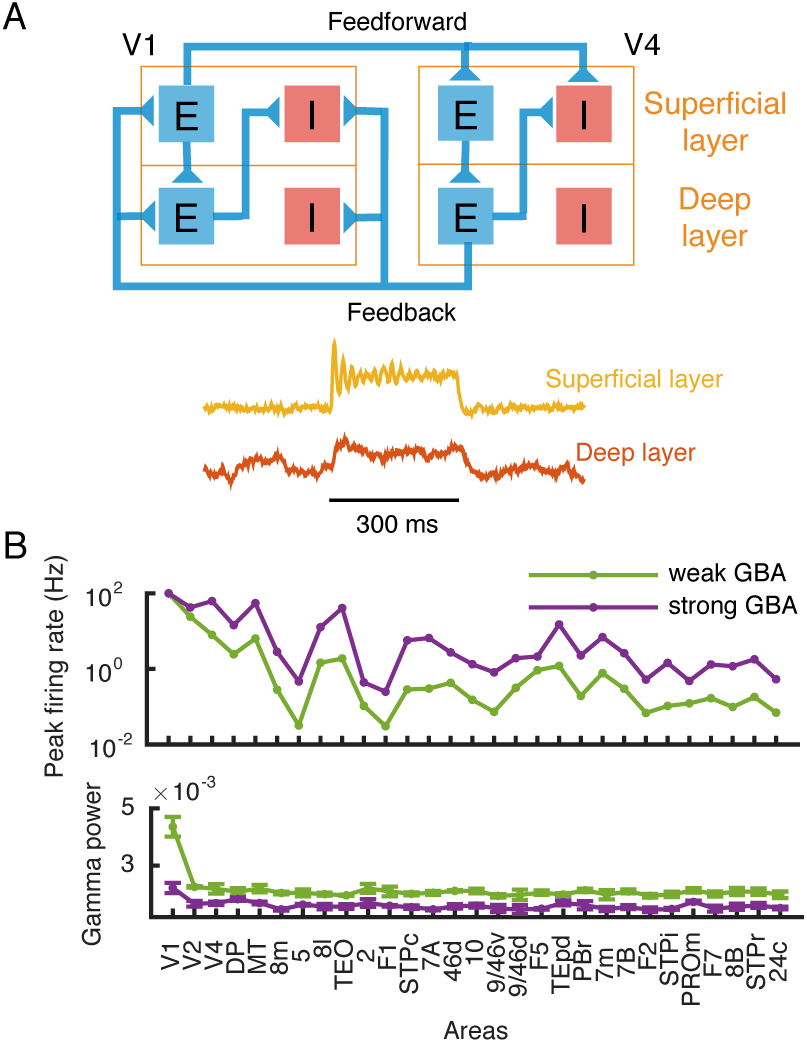
Increasing Global Balanced Amplification also improves signal propagation in a large-scale model endowed with a laminar structure. (A) (Upper) Circuit diagram showing the intra and inter-areal connectivity in the large-scale model with a superficial layer and a deep layer in each local area. In addition to the connections shown, the E-I circuit in each layer of every area has local connectivity as in Fig. 1A. (Lower) Sample oscillatory activity for the local-circuit in the superficial layer (top) and deep layer (bottom), in response to a brief input of 300 ms given to the excitatory population in the superficial layer of V1. (B) (Upper) Peak firing rate across areas as the response to a pulse input to V1 propagates along successive areas in the hierarchy, for weak (green) and strong (purple) GBA. (Lower) Gamma power across cortical areas (except for early areas) is not dramatically changed from the weak (green) to strong (purple) GBA regime.

### Two (asynchronous and synchronous) regimes of transmission in a large-scale spiking network model

Signal transmission in neural systems is mediated through spiking activity, therefore it is crucial to assess our mechanism in a more realistic spiking neural network model. We thereby extend our investigation by building a large-scale spiking network model (Fig. 4A, see Methods). Inter-areal connectivity is based on the same anatomical data used above [9], and the inter-areal delays are introduced by considering the corresponding interareal wiring distances [9] and assuming a constant axonal conduction velocity. Propagation of spiking activity has been previously studied in the context of asynchronous firing rates [3, 4, 18] and synchronous activity [5–7], both of which have been observed experimentally [32–35].

**Fig. 4:**
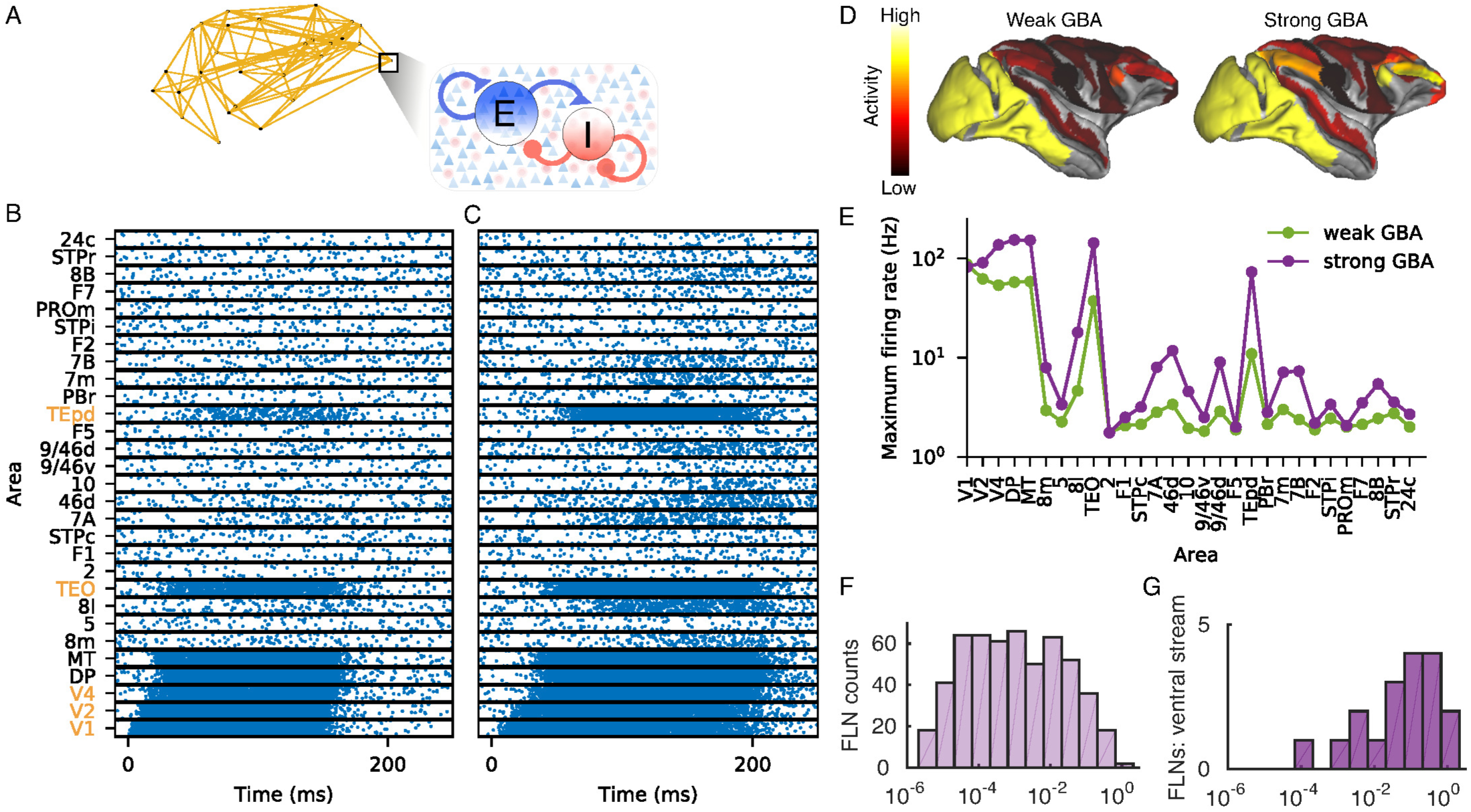
Reliable signal propagation in the asynchronous regime in a spiking network model. (A) Circuit diagram for local-circuit connectivity for the spiking network model. Each cortical area contains 1600 excitatory and 400 inhibitory neurons. (B) Response to a 150 ms pulse input to V1, as it propagates along the hierarchy. The areas along the ventral stream showing strong response activity are indicated with orange labels. (C) Similar to (B), but with strong GBA. (D) Spatial activity pattern across the macaque cortical surface corresponding to parameters in (B) and (C). (E) Peak firing rate across areas as the response to a pulse input to V1 propagates along the hierarchy for weak (green) and strong (purple) GBA. (F) FLN strengths of long-range connectivity across areas span five orders of magnitude. (G) FLN strengths along the ventral visual stream areas are stronger than the average, leading to more effective signal propagation along the ventral pathway than the dorsal pathway.

We first evaluate the performance of our mechanism in the asynchronous regime (Fig. 4), and later in the synchronous regime (Fig. 5). To examine asynchronous propagation, we stimulate V1 with a long (150 ms) pulse input. The corresponding raster plot for weak GBA (Fig. 4B) shows a strong response activity only in early visual areas. Weak activity is observed in a part of the frontal eye fields (area 8l), but is conspicuously absent from the dorsolateral prefrontal cortex (dlPFC), which has been associated with working memory and decision making [36]. Increasing GBA facilitates signal propagation to higher cortical areas (Fig. 4C). Stronger response activity is observed in the higher areas including those in the dLFPC (area 46d, 9/46d), the frontopolar cortex (area 10), parietal area 7 (7A, 7B, 7m) in the dorsal stream, and in the frontal eye fields (area 8l, 8m), as indicated by the peak firing rate responses across areas (Fig. 4D,E). For weak GBA, signals propagate mainly along the ventral visual stream (V1, V2, V4, TEO and TEpd) (Fig. 4B). This is presumably due to the much stronger anatomical projection weights between these areas compared to the overall connectivity (with a significant difference between their average projection strengths, P = 0.015) (Figs. 4F,G). These stronger weights enable propagation along ventral areas through recurrent excitatory loops, while the signal fails to reach higher areas due to relatively weak connections.

**Fig. 5:**
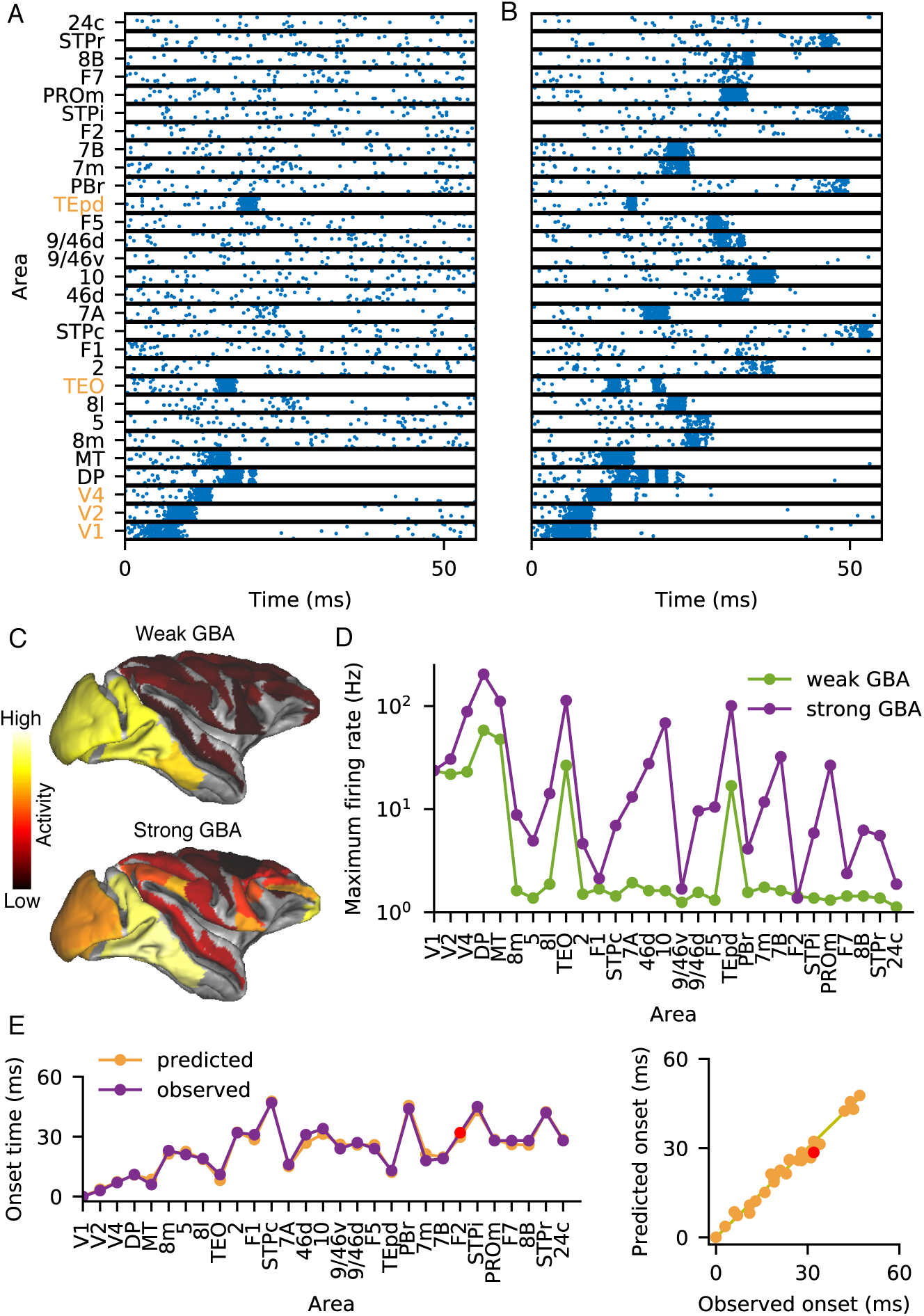
Reliable signal propagation in the synchronous regime in a spiking network model. (A) Response to a brief input to V1 as it propagates along the hierarchy. The areas along the ventral stream showing strong response activity are indicated with orange labels. (B) Similar to (A) but with strong GBA. (C) Spatial activity pattern across the macaque cortical surface corresponding to parameters in (B) and (C). (D) Peak firing rate across areas as the response to the pulse input to V1 propagates along the hierarchy for weak (green) and strong (purple) GBA. (E) (Left) Predicted and observed response onset times across areas as the signal propagates along the hierarchy. Area F2 which fails to show response activity is indicated in red. (Right) Predicted and observed onset times from (B) where each dot indicates an area. The line indicates the diagonal *y* = *x*.

Even with strong GBA, many areas do not show noticeable responses to an input to V1. These previously silent areas can however be activated when the input is directed to a different sensory area. For example, an input to the primary somatosensory cortex (area 2) uncovers a new set of areas showing propagation (Fig. S5), with the same connectivity parameter values used in Figs. 4B,C. Propagation was observed largely in the somatosensory areas of the parietal lobe, and extended to prefrontal areas for strong GBA.

Our model displays a second mode of transmission, in the synchronous regime. Following previous works on synchronous propagation [5, 37], we stimulated V1 with a brief (10 ms) input pulse rather than a long-lasting stimulus. Connectivity is set stronger as compared to the asynchronous propagation case (see Methods) to allow for a quick build-up of network activity, since stronger connectivity leads to higher degree of population synchrony (Fig. S6). For weak GBA, signal propagates in the visual areas, but it does not reach higher cognitive areas in prefrontal cortex (Fig. 5A), as in the case of the asynchronous model (Fig. 4B). Increasing GBA enables successful signal propagation to several higher areas including those in the dlPFC (areas 46d, 9/46d) and the frontal eye fields (areas 8l, 8m) (Fig. 5B,C,D).

Early response onset occurs along the ventral stream (Fig. 5B); the signal then propagates to higher cognitive areas, and eventually to the superior temporal polysensory (STP) areas involved in multisensory integration, which form part of a cluster that shapes functional connectivity [15]. After testing multiple hypotheses (Fig. S7), we found that the observed response onset times in our large-scale network are best predicted by a shortest-path toy model (Fig. 5E). In this toy model, we first ignore projections whose strength falls below a certain threshold value, and then assume that the signal, starting from V1, follows the shortest possible path to reach any given area. The shortest path is determined based on the anatomical inter-areal wiring distances [9], and a constant conduction velocity is assumed (see Methods). While our results provide reasonable response onset latencies in general, the values found for dorsal stream visual areas are somehow smaller in our model than those observed in anesthetized macaques [38]. This could be attributed to the absence of dorsal areas V3 and MST in our anatomical subgraph [9], and the lack of separate pathways, for example magnocellular and parvocellular pathways, in our modeled areas.

### Threshold-crossing for signal propagation and conscious perception

The emergence of activity across several cortical areas in Figs. 4C, 5B is reminiscent of the “global ignition” observed during conscious perception [20, 21, 39, 40]. Global ignition is characterized by a distributed cerebral activation pattern, contingent on strong parieto-frontal network activation that emerges when the bottom-up input exceeds a certain threshold [20]. To examine these phenomena, we monitor the activity across different cortical lobes in our asynchronous propagation model (Fig. 4C) on successively increasing the input current strength arriving at V1. From our original simulations on the large-scale spiking model with improved propagation (Fig. 4C,E), we estimate that the signal from V1 reaches around 16 cortical areas, which here we will refer to as “active areas” (see Methods). On varying the input strength, the normalized peak response across these areas (Fig. 6A) reveals a clear separation across the four cortical lobes. Low input (Fig. 6A) shows activation only in the early visual areas in the occipital lobe, similar to subliminal processing [20] associated with weak stimulus strength. As we increase the input, activity starts to emerge in the temporal lobe including areas in the ventral stream, followed by parietal activation including area 7 involved in visuo-motor coordination. Activity across areas in the prefrontal cortex, a necessary requirement for conscious processing [20], emerges at similarly high values of input current [20] (Fig. 6A). Simultaneous emergence [41] is observed in the densely connected hub of the prefrontal cortex [14], including the dlPFC, frontopolar cortex and the frontal eye fields (Fig. 6A).

**Fig. 6:**
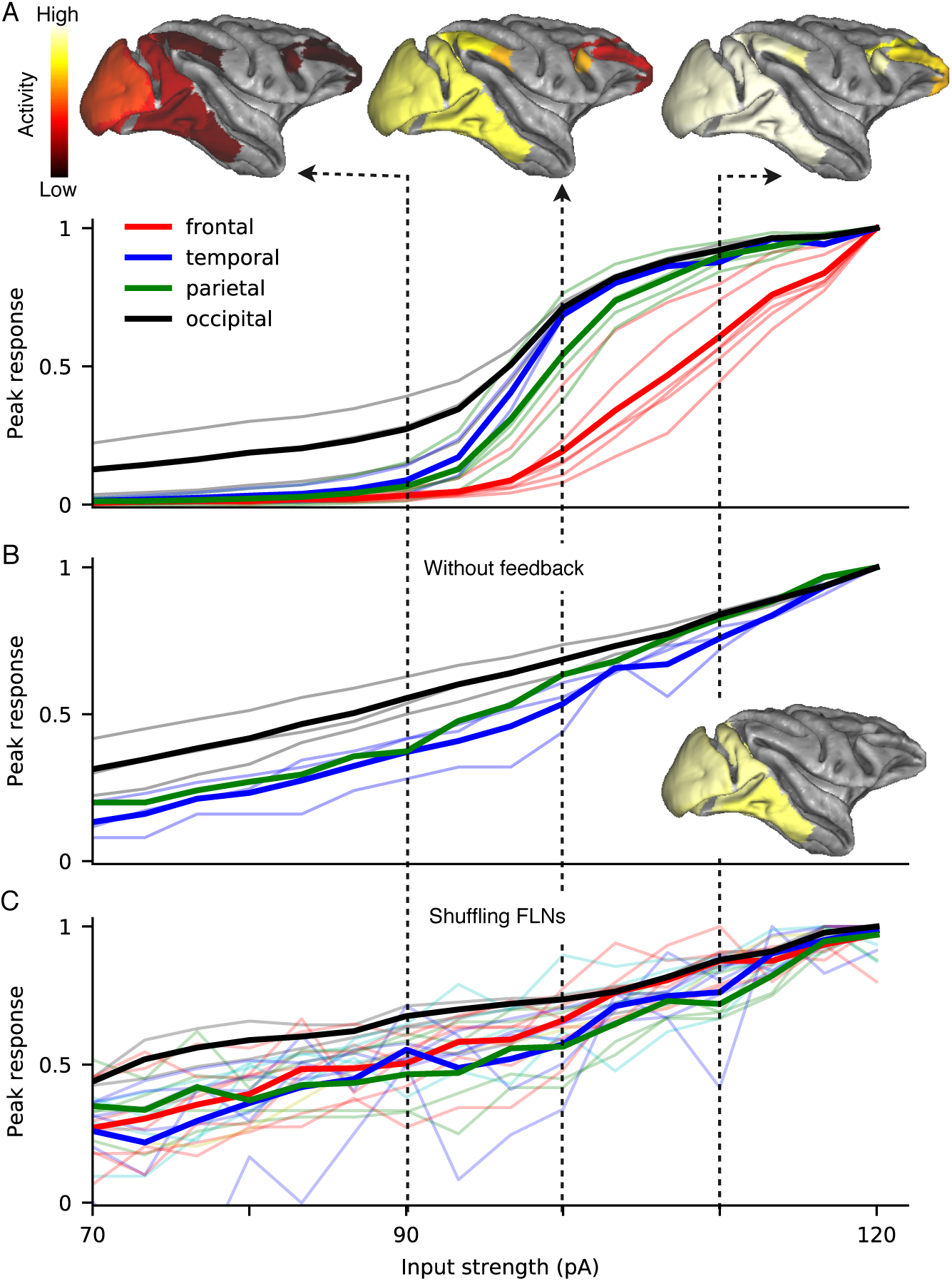
Threshold-crossing of input strength is required to engage the parietal and frontal lobes, in support of the global ignition model of consciousness. (A) (Lower) Normalized peak population activity in the active areas across cortical lobes, as a function the input current strength, using the parameters corresponding to Fig. 4C. The light colored lines correspond to individual areas. The thick lines show the mean activity across all the active areas in each cortical lobe, revealing that activation of the temporal lobe (blue), parietal lobe (green) and frontal lobe (red) requires threshold-crossing of input strength. (Upper) Spatial activity (indicated by color) across the macaque cortical surface, for the current strength of 90, 100, 110 pA respectively. (B) (Left) Similar to (A) with the deletion of all feedback projections in the model. (Inset) Spatial activity pattern of the macaque cortical surface at input strength of 110 pA, to be compared with the intact model (top right in (A)). (C) Similar to (A) with scrambling the long-range inter-areal connectivity while maintaining the network topology. In both (B) and (C) the input threshold-crossing for activation of association areas is no longer present, indicating the importance of the inter-areal cortical connectomics of macaque monkey including feedback loops for this phenomenon.

When feedback projections are removed from the large-scale network (Fig. 6B), we observe that the signal fails to reach prefrontal areas even for strong input current. Activity becomes restricted largely to the occipitotemporal region, in the early visual areas and along the ventral stream. This activity pattern in the absence of feedback projections is similar to that of preconscious processing, associated with the absence of top-down attention [20].

Finally, we tested whether the quantitative anatomical connectivity structure has a key role in determining the activity pattern across the cortical lobes observed in our model (Fig. 6A). To do so, we randomly rewired the anatomical projection strengths (only for those projections with non-zero connection weights) while maintaining the network topology. With scrambled quantitative connectivity, the response curve as a function of input strength becomes similar across the temporal, parietal and frontal lobes (Fig. 6C) and the threshold-effect disappears. This finding indicates that the quantitative connectivity structure has a critical role in the emergence of lobe-specific activity and the higher input threshold for prefrontal activity; the heterogeneity across areas and the network topology are not enough, by themselves, to explain these phenomena.

## Discussion

Transmission of signals is essential for neural coding of external stimuli, and represents a major topic about the dynamical operation of a multi-regional cortical system [1, 2]. However, most previous models for signal propagation used a purely feedforward network architecture, in contrast to the recent report that about half of inter-areal cortical connectivity consists of feedback projections [9] (Fig. S1). We re-examine the long-standing problem of signal transmission in large-scale circuit models that incorporate heterogeneity across areas and mesoscopic inter-areal connectivity data [9]. The central idea of this paper is a generalized balanced amplification mechanism in which strong long-range excitatory coupling produces a transient amplification of signals, balanced by enhanced local inhibitory-to-excitatory strength to ensure network stability. Generalized balanced amplification provides a solution to the tradeoff between the need of sufficiently strong excitation for reliable signal transmission and the risk that inter-areal recurrent excitation potentially destabilizes the entire system. We found that this mechanism improves signal propagation by up to 100 fold in large-scale network models of population firing rates with [16] and without [15] a cortical laminar structure, as well as a network model of spiking neurons. Furthermore, inter-areal connection strengths along the ventral visual stream areas are stronger than the overall long-distance connection strengths, which underlies more effective signal propagation along the ventral pathway than the dorsal pathway, a prediction that can be tested experimentally. Finally, surprisingly, our model reproduces signature dynamics of "global ignition" associated with conscious report of a sensory stimulus.

Propagation of spiking activity in feedforward networks has been extensively examined theoretically, in terms of the propagation of both asynchronous [3, 4, 18, 42], and synchronous [5–7] spiking activity. Synchronous propagation in a feedforward chain can be analyzed in detail, in terms of temporal jitter [8], refractoriness [43], and the distribution of synaptic strengths [44]. In addition, feedback from a higher area to the inhibitory interneurons in a lower area has been proposed to improve synchronous propagation in a multilayer network [37]. From a more general point of view, Kumar and collaborators [7] studied synchronous and asynchronous propagation in a feedforward network embedded in a recurrent network. Gating mechanisms to control propagation of asynchronous [18] and synchronous [45] activity have been proposed, based on the balance [18] and latency [45] between excitation and inhibition, and on other biophysical properties [46]. The asynchronous and synchronous modes of propagation have been integrated in a recent study on feedforward propagation [2], which argues that the two modes represent two extremes of a continuum parametrized by the model parameters. While previous theoretical works largely investigate either synchronous or asynchronous propagation, our propagation mechanism is effective for both the modes (Figs. 4, 5).

Another important insight from this work is to elucidate how signal propagation depends on the degree of non-normality of the underlying network connectivity. Signals in a dynamical system by a normal connection matrix will inevitably attenuate or grow exponentially leading to instability. However, as pointed out in previous work [17], connection matrices of biological neural circuits are always non-normal due to the Dale’s law. When connectivity is governed by strong recurrent excitation balanced by strong inhibition, biological circuits can transiently amplify incoming signals. This phenomenon, termed as balanced amplification [17], is proposed to ubiquitously contribute to neural dynamics across the brain [17]. For example, balanced amplification has been recently shown to improve memory replay through signal amplification in a hippocampus model [47]. Our work extends this basic dynamical motif of balanced amplification from a local circuit to a multi-regional large-scale system to explain reliable cortical signal transmission.

### Future extension of our cortical circuit model

In this work simulations have been limited to one sensory modality at a time. This can be easily extended by considering multiple input modalities simultaneously presented to the model, to study how signals from different sensory modalities may interact and aid one another. Other mechanisms besides GBA that improve propagation should also be examined. For example, increasing the local recurrent inhibition and balancing it with local excitatory-to-inhibitory strength is expected to have a similar effect. This is because an increase in the excitatory-to-inhibitory connection strength would have a stronger effect along the hierarchy following from the scaling of excitatory projections. This would consequently scale the suppression of inhibition, resulting in a higher disinhibition along the hierarchy and ultimately to a relative increase of excitation in higher cortical areas.

While the anatomical projection strengths used in our model [9] span five orders of magnitude (Fig. S1), the role of weak connections in propagation remains unclear. For instance, in a phenomenological network model using the same dataset [25], the authors found that their measure of information transfer was robust to the removal of a significant fraction of the weakest connections, and depended largely on the stronger connections. Similarly, we observe a minor change in the propagation ratio after removing the weakest connections in the rate model (Fig. S3), even as the network density is reduced to 50% of its original value. We also observe that the response latencies for synchronous propagation (Fig. 5B) can be predicted by thresholding the anatomical connection weights to include only the strongest projections (Fig. 5E). Weaker inter-areal projections can, however, be functionally important if they target a small neural population that has a strong impact in the local circuit. More data on cell-type specific cortical connectivity is needed to shed light on this issue.

### Role of subcortical structures

Although our model takes into account cortical connectivity data, which forms a large part of the input to cortical areas [10], the role of subcortical structures remains to be explored, and emerging subcortical connectivity data should be incorporated. For instance, cortico-thalamo-cortical interactions have been shown to drive robust activity in the higher-order somatosensory cortex [48]. Experimental studies have also shown that the pulvinar synchronizes activity [49] between interconnected cortical areas, indicating its role in regulating information transmission across the visual cortex. Recent work [42] studies asynchronous cortical transmission through two multilayered feedforward networks, and explores the role of long-range inhibitory pulvinar connections in linking multiple cortical stages to boost propagation. While our large-scale models only employ anatomical connectivity from macaque cortex, novel thalamocortical connectivity data from the mouse [12] could be used to examine our mechanism in a corticothalamic large-scale model of the mouse brain.

### Ignition theory

Recent work suggests that the emergence of parieto-frontal activity could be viewed as a precursor to conscious perception [41]. Such an event would follow from the input exceeding a threshold, leading to a reverberating neuronal assembly [20]. In this sense, subliminal, preconscious and conscious processing would be associated with different levels of top-down attention and bottom-up stimulus strength [20]. Our large-scale model, able to propagate signals efficiently (Fig. 4), can be used as a framework for a computational examination of these phenomena. Subliminal processing, characterized by weak bottom-up activation insufficient to trigger large-scale reverberation [20], resembles the weak input case in Fig. 6A. Preconscious processing is characterized by the absence of top-down attention [20]. Correspondingly, we observe a disruption of global reverberation after removing top-down projections (Fig. 6B) that mostly originate from the strongly connected core of prefrontal and association areas [14]. Conscious perception requires both strong bottom-up stimulus and top-down attention [20]. In the intact network model, strong input elicits activity across several prefrontal and parietal areas (Fig. 6A), which can be viewed as a necessary condition for conscious perception. Although a complete characterization of the observed neural activity related to conscious perception is beyond the scope of this work, we believe that future extensions of our network model would constitute a powerful tool to understand this phenomenon. In particular, NMDA receptors may be more prominent at synapses of top-down projections than those of bottom-up projections [50]. Implementing NMDA dynamics in the spiking network model could provide insight on the nature of the top-down amplification of posterior areas following prefrontal activation, as observed in conscious perception tasks [20].

We are ushering in a new era of understanding large-scale brain systems beyond local circuits. Whereas brain connectomics is essential, structural connectivity is insufficient to predict dynamical behavior of recurrent neural circuits. Our work on reliable signal propagation offers another demonstration of this principle, and represents an important step in our investigations of cognitive processes in a large-scale brain circuit.

## Supplemental Information

Supplemental Information includes 7 Supplementary figures, and an Appendix.

## Author Contributions

M.R.J., J.F.M., G.R.Y. and X.-J.W. designed the research, had regular discussions throughout the project and wrote the manuscript. M.R.J. performed the computational research.

## Acknowledgements

Funding was provided by The Swartz foundation, ONR Grant no. N00014-17-1-2041 and Simons Collaborative Global Brain (SCGB) Program Grant to X.-J.W‥

## Methods The rate model

The model examined in Figs. 1, 2, based on a recently published large-scale model of the macaque cortex [15], is described below. The dynamics of each area is described as a threshold linear recurrent network, with interacting excitatory and inhibitory populations, as follows

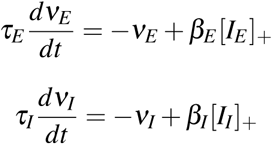

where [*I*_*E*_]_+_ = max(*I*_*E*_,0). *ν*_*E*_, *ν*_*I*_ denote the firing rates of the excitatory and inhibitory populations respectively, and *τ*_*E*_, *τ*_*I*_ are the corresponding intrinsic time constants. *β*_*E*_,*β*_*I*_ are the slopes of the f-I curves. The local-microcircuit is qualitatively the same across areas, with quantitative inter-areal differences as stated below.

### Heterogeneity across areas

The laminar pattern of inter-areal projections is used to assign a hierarchical position to each area [15, 23, 51]. This is based on the notion that feedforward projections tend to originate from the superficial cortical layer, and feedback projections from the deep layer [28]. Thus the hierarchical distance between a source and target area is computed based on the fraction of projections originating in the superficial layer of the source area [15]. An area’s hierarchical position is found to be strongly correlated with the number of basal-dendritic spines of layer 3 pyramidal neurons in that area [15, 22]. The pyramidal-cell spine count increases with the hierarchical position of the cortical area by a factor of 6~7 [15, 22], and it is used as a proxy for the total excitatory drive. Thereby, heterogeneity across areas is introduced in the form of a gradient of excitatory connection strengths along the hierarchy [15].

Inter-areal projection strength is based on a recently published anatomical connectivity dataset from the macaque cortex [9]. The data measures the number of neurons labelled by a retrograde tracer injected in 29 widely distributed cortical areas. To control for the injection size, the neuron counts are normalized by the net number of neurons labelled by the injection, resulting in an FLN (Fraction of Labelled Neurons) across two areas. Thus, given areas *i*, *j*, the *FLN*_*ij*_ is the number of neurons projecting from area *j* to area *i* weighted by the net number of neurons projecting to area i from all the areas. The net incoming current is given by

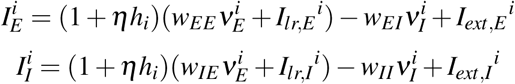

Where 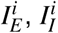 denote the input currents to the excitatory and inhibitory populations respectively for area *i* and *w*_*ij*_ denotes the local-circuit connection strength from the population type *j* to the population type *i*. 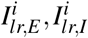 correspond to the long-range input currents, assumed to be purely excitatory. 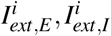 correspond to the external inputs. The hierarchical position *h*_*i*_ is normalized to lie between 0 and 1. *η* scales the excitatory connection strengths based on the hierarchical position of the area. We set *η* to 0.68 [15]. The background firing rate, is set to an excitatory rate of 10 Hz, and an inhibitory rate of 35 Hz [15]. The background rate is subtracted when monitoring the response activity in Figs. 1, 2 and Figs. S2, S3. The parameter values are set as *τ*_*E*_ = 20, *τ*_*I*_ = 10 (in ms), *β*_*E*_ = 0.066, *β*_*I*_ = 0.351 [15, 52]. The long-range input currents are given by,

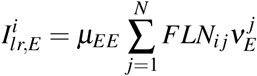

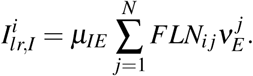

Thus, the inter-areal connectivity depends on the corresponding FLNs and is scaled by the global scaling parameters *μ*_*EE*_ and *μ*_*IE*_ corresponding to long-range *E* to *E* and long-range *E* to *I* coupling respectively. The connectivity parameters are set as *μ*_*IE*_ = 25.3, *w*_*EE*_ = 24.3, *w*_*IE*_ = 12.2, *w*_*II*_ = 12.5 (in pA/Hz) [15]. The connectivity parameters corresponding to the local I to E connection strength and the global excitatory coupling are (in pA/Hz) *w*_*EI*_ = 19.7, *μ*_*EE*_ = 33.7 for weak GBA and *w*_*EI*_ = 25.2, *μ*_*EE*_ = 51.5 for strong GBA (Fig. 2).

### Local balanced amplification

For Fig. 2B, recurrent excitation and lateral inhibition values are (6, 6.7) and (4.45, 4.7) for strong LBA and weak LBA respectively. The excitatory to inhibitory connection strength, and the inhibitory recurrent strength values are fixed at 4.29, 4.71 respectively (Fig. 2B, C).

### The laminar model

The model examined in Fig. 3 is based on a recently published large-scale rate model of the macaque cortex [16], which incorporates a cortical laminar structure. The intra-laminar cortical circuit in each area consists of a recurrently connected excitatory and inhibitory population, with dynamics described by the following Wilson-Cowan equations,

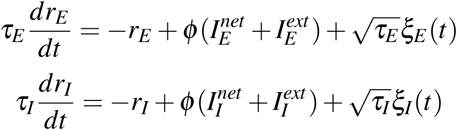

where *r*_*E,I*_ denote the dimensionless mean firing rates of the excitatory and inhibitory populations respectively, *τ*_*E,I*_ denote the corresponding time constants, *ξ*_*E*,*I*_ denote Gaussian white noise terms with strengths *σ*_*ε,I*_, and *ϕ*(*x*) = *x*/(1 – exp(–*x*)) is the transduction function. The network input, denoted by 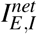, is the input arriving to the *E*, *I* populations respectively from the other populations in the network, and includes the inputs from the same layer, a different layer, and from different areas. The external input, denoted by 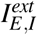, is the input arriving from external sources such as sensory stimuli, thalamic input and other cortical areas not explicitly included in the model. The network input taking into account only local contributions, that is, on assuming an isolated intra-laminar population, is given by,

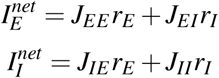

where *J*_αβ_ denotes the mean synaptic strength from population *β* to population *α*. The parameter values for the circuit in the superficial layer are *τ*_*E*_ = 6 ms, *τ*_*I*_ = 15 ms, *J*_*EE*_ = 1.5, *J*_*IE*_ = 3.5, *J*_*EI*_ = –3.25, *J*_*II*_ = –2.5 and *σ*_*E*,*I*_ = 0.3. The parameters for the circuit in the deep layer are the same except for *τ*_*E*_ = 30 ms, *τ*_*I*_ = 75 ms, and *σ*_*E*,*I*_ = 0.45.

The inter-laminar interactions assume only the strongest connections between the superficial and deep layer, that is, the excitatory projections from the pyramidal neurons of the superficial layer to the pyramidal neurons of the deep layer, and those from the pyramidal neurons of the deep layer to the interneurons of the superficial layer. The long-range connections are assumed to be excitatory. The inter-areal interactions assume that feedforward projections originate from the superficial layer and target the excitatory population in the superficial layer across areas. Feedback projections, are assumed to originate from the deep layer and target both the E and I populations in the superficial and deep layer across areas. The feedforward connections from the excitatory neurons of the superficial layer to the inhibitory ones (Fig. 3A) were added to test our mechanism and are not present in the original model [16]. These connections are assumed to have 20% of the strength of the feedforward connections targeting the excitatory populations across areas. For consistency with the other models examined in the present work, we simulate the large-scale laminar model considering 29 areas as opposed to the 30 areas [16] (i.e. we remove area LIP).

### The spiking network model

We build a spiking network model examined in Figs. 4, 5, 6 Simulations are performed using a network of leaky integrate-and-fire neurons, with the local-circuit and long-range connectivity structure similar to the rate model. Each of the 29 areas consists of 2000 neurons, with 1600 excitatory and 400 inhibitory neurons. Connection density, both intra and inter-areal is set at 10%. The membrane constant values are *τ*_*E*_ = 20 ms for excitatory, and *τ*_*I*_ = 10 ms for inhibitory neurons. The resting membrane potential *V*_*r*_, reset potential *V*_*reset*_, and threshold potential *V*_*t*_ are given by *V*_*r*_ = ‐70 mV, *V*_*reset*_ = –60 mV and *V*_*t*_ = ‐50 mV respectively, and the absolute refractory period *τ*_*ref*_ = 2 ms. Background currents are injected to yield firing rates in the 0.75-1.5 Hz range for the excitatory, and 5-6 Hz for the inhibitory population in the absence of input. We introduce distance-dependent inter-areal synaptic delays by assuming a conduction velocity of 3.5 m/sec [16, 53] and using a distance matrix based on experimentally measured wiring distances across areas [9]. Inter-areal delays are assumed to have a gaussian distribution with mean based on the inter-areal wiring distance and variance given by 10% of the mean. Intraareal delays are set to 2 ms. For neuron *i*, the depolarization voltage at the soma follows

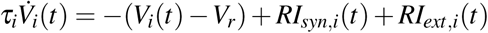

where *I*_*syn*,*i*_(*t*) is the post-synaptic current and *I*_*ext*,*i*_(*t*) is the external input. The post-synaptic current corresponds to a summation of spike-contributions of spikes arriving at different synapses at different time intervals, where the spikes are modeled as delta functions [19]. Thus,

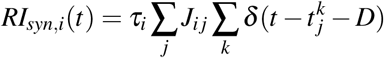

where *τ*_*i*_ is the membrane constant, *D* is the transmission delay, 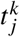 is the emission time of the *k*^*th*^ spike at the *j*^*th*^ synapse, and *J*_*ij*_ is the synaptic strength of the *j*^*th*^ synapse to neuron *i*. We choose *R* = 50*MΩ*; synaptic strengths for the local and global coupling parameters are chosen in the range 0.01 ‐1 mV [54, 55]. Simulations are performed using the Python library Brian2, using a time-step of 0.1ms. Population firing rate in each case is calculated using sliding time window with a bin size of 10 ms and the sliding window width of 1 ms. For the spiking model simulations, the parameter *η*, governing the gradient of excitatory strengths, is set to 4.

For the asynchronous regime (Fig. 4), the synaptic strengths are set to (in mV) *w*_*EE*_, *w*_*IE*_, *w*_*II*_, *μ*_*IE*_ = 0.01, 0.075, 0.075, 0.19/4. The global excitatory coupling and local I to E strength are set to (in mV) *μ*_*EE*_, *w*_*EI*_ = 0.0375,0.0375 for weak GBA, and (in mV) *μ*_*EE*_,*w*_*EI*_ = 0.05,0.05 for strong GBA (Fig. 4). The input current duration is set to 150 ms, based on recent work suggesting that activity packets of 50-200 ms duration serve as basic building blocks of global cortical communication [56]. The input current strength is set to 300 pA in Fig. 4B and 126 pA in Fig. 4C resulting in a peak firing rate of 82-87 Hz in V1. For weak and strong GBA in Fig. S5, connectivity parameters were used as in Fig. 4B, C. A pulse input of 150 ms duration is injected to the excitatory population of area 2. The input current strength is set to 138 pA for weak GBA and 140 pA for strong GBA, each resulting in a peak firing rate of ~40-41 Hz in area 2.

For the synchronous regime in the spiking model, we set the synaptic strengths *w*_*EE*_, *w*_*IE*_, *w*_*II*_,*μ*_*IE*_ to be a multiple of the values used in the asynchronous case, the values are (in mV) *w*_*EE*_, *w*_*IE*_, *w*_*II*_,*μ*_*IE*_ = 0.04, 0.3, 0.3, 0.19. The global excitatory coupling and local I to E strength are set to (in mV) *μ*_*EE*_, *w*_*EI*_ = 0.16,0.56 for weak GBA, and *μ*_*EE*_, *w*_*EI*_ = 0.25,0.98 for strong GBA (Figs. 5A, B). The input current duration is set to 8 ms, and the input current strength is set to 200 pA, resulting in a peak firing rate of ~24 Hz in V1. For Fig. 4D, we map the attenuation in Fig. 4E logarithmically to a heatmap. For better visualization, we threshold the areas showing strong propagation and zoom in on the other areas, and plot the values using Caret [57]. For Fig. 5C, we map the attenuation in Fig. 5D logarithmically to a heatmap and plot the values using Caret [57].

### Predicted onset times

The response onset time for a given area in Fig. 5B is defined as the time at which the response activity starts building up in that area prior to the peaking of activity. To compute the predicted onset times (Fig. 5E), the FLNs are thresholded to 0.02, that is, connections with FLN values below 0.02 are removed, following which the predicted onset time is computed assuming that the signal follows the shortest possible path to reach a given area. For a positive integer *k*, a *k*-step path, given by {*A* = *A*_0_,*A*_1_,*A*_2_,…,*A*_*k*_ = *B*} is said to exist between areas *A* and *B* if there exist anatomical connections from area *A*_*m*_ to area *A*_*m*+1_ from *m* = 0,1,…, *k* – 1, that is, if *FLN*_*A*_*m*+1_,*A*_*m*__ > 0. We use Dijkstra’s algorithm to compute the shortest path from V1 to a given area based on the inter-areal wiring distances [9]; the predicted onset time is computed assuming a constant conduction velocity of 3.5 m/s [16, 53]. Let *S*_*A*,*V*1_ be the length of the shortest path from *V*1 to a given area *A*, divided by the conduction velocity. We assume that the signal undergoes local-circuit processing at each step while traveling from V1 to area *A*, and assume a processing time of 1 ms in each area. (This is also assumed in Supplementary Fig. 5.) Assuming that the shortest path from V1 to area *A* is a *k*-step path, the predicted onset time (in ms) for the signal to reach area *A* from V1 is given by *S*_*A,V*1_ + *k*. While computing the predicted onset times, we eliminate those areas from our computations wherein the signal does not elicit response activity. In particular, as the background firing rate for the excitatory population lies in the range 0.75-1.5 Hz, we eliminate those areas with a peak firing rate < 1.5 Hz. The only such area we find is F2 (in the premotor region). For F2, the predicted onset time for visualization purposes (Fig. 5E) is set as the mean of the predicted onset times corresponding to its neighboring areas in the hierarchy.

### From the asynchronous regime to the synchronous regime in the parameter space

The degree of population synchrony *χ*, in the network is measured based on the variance of the average population voltage in comparison to the variance of the individual neuron voltage. We compute *χ*, which takes a value between 0 and 1, defined [58] as

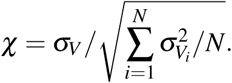

*χ* is computed (Fig. S6) on moving from the parameter set in Fig. 4B to the parameter set in Fig. 5A. The parameter set comprises of the inter-areal and intra-areal connectivity parameters, the current duration and input current strength. The current duration corresponding to the parameter set in Fig. 5A is set to 100 ms to characterize the fluctuations in the average population voltage. For each parameter set, the simulation is run using 5 random seeds, to get the mean value and standard deviation of *χ*.

### Population activity across cortical lobes with varying input strength

Active areas in Fig. 6A are defined as those for which the peak firing rate in Fig. 4C is at least 5% that of the corresponding peak rate in V1. The population activity of these areas is monitored for the input current strength varying from 70 to 120 pA. At each current value, we subtract the background firing rate for each of the active areas. To compare activation across areas, the normalized activity for each active area is computed. The normalized peak firing rate for a given current value is computed by dividing the peak firing rate at that value by the maximum peak firing rate over all the current values, so that the normalized rate lies between 0 and 1 as the current strength is varied. Active areas in Fig. 6A are occipital areas V1, V2, V4, parietal areas DP, 7A, 7m and 7B, temporal areas MT, TEO and TEpd and prefrontal areas 8m, 8l, 46d, 10, 9/46d and 8B. In the absence of feedback in Fig. 6B, normalized activity is examined only in those areas deemed “active”, for which the peak firing rate at current strength 120 pA is at least 1% that of the peak firing rate in V1. For Fig. 6C, on scrambling the anatomical connectivity, the areas which show a strongly non-monotonic change in normalized activity in response to an increase in current strength, are regarded as not receiving the input signal, and are not plotted. For each input current value in Fig. 6, the simulation is run using 5 random seeds; the curves plotted for each area correspond to the mean from the 5 simulations. For Figs. 6A (upper) and B (inset), we map the difference between the normalized activity of an area and 1, for the given current value, to a heatmap and plot using Caret [57].

## Supplementary Information

Fig. S1 shows the distribution of the anatomical FLN weights, overlaid with the feedback FLNs, which comprise ~50% of the total connections, and overlaid with the unidirectional FLNs, which comprise ~20% of the projections.

Fig. S2 tests the robustness of the mechanism for the rate model (Fig. 2) under several conditions; the mechanism continues to yield strong improvement in propagation across these conditions. The mechanism is tested in the absence of feedback connections (Fig. S2A), and in the absence of heterogeneity across areas, that is, on setting the parameter η, which governs the gradient in excitatory connection strengths, to 0 (Fig. S2B). Removing feedback (Fig. S2A) worsens propagation particularly for strong GBA with high inter-areal coupling.

We test our mechanism after symmetrizing the anatomical connectivity matrix, based on the geometric means (Fig. S2C) and the arithmetic means (Fig. S2D) of the underlying inter-areal projection strengths. We find that the improvement in propagation doesn’t depend on the specific directionality of connectivity, which is not revealed by the commonly used method of diffusion tensor imaging. Symmetrizing the FLN matrix using the geometric means weakens propagation, which can be attributed to the removal of one-directional connections; the data reveals ~20% one-directional connections (Fig. S1).

To investigate a correspondence with the “small-world” property of biological networks of high clustering and short path lengths, Fig. S3 tests the mechanism on removing the weak anatomical connections with strengths below a parametric threshold [25]. Varying the threshold reveals that the mechanism consistently shows a strong improvement in propagation even as the network density is reduced to 50% of its original value. A small change in the propagation ratio is observed on thresholding the FLNs, even as a large fraction of the weakest connections are removed.

Since inhibition-stabilized networks showing balanced amplification are known to be characterized by strong non-normality of the underlying connectivity matrix [17], we examine the non-normality of the large-scale connectivity matrix on increasing GBA (Fig. S4) corresponding to the parameters used in Fig. 2I. Non-normality can be examined through Schur decomposition [17, 26], which can be used to express the effective connectivity across network basis patterns through self-connections and feedforward connections. For a given matrix *A*, the Schur decomposition of *A* can be expressed as, *A* = *UTU*\* where *U* is a unitary matrix, and *T* is an upper triangular matrix such that *T* = Σ + *R*, where Σ is a diagonal matrix and *R* is strictly upper triangular. Based on the non-normality measure by Henrici [26], the departure of *A* from normality can be approximated as

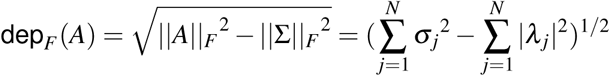

where *σ*_*j*_, *λ*_*j*_, denotes the singular value and the eigenvalue of *A* respectively, for a given *j* and || ||_*F*_ denotes the Frobenius norm. For normal matrices, *σ*_*j*_ = |*λ*_*j*_| for each *j*, thus dep_*F*_(*A*) = 0. An improvement in propagation is accompanied by an increase in the non-normality measure of the underlying connectivity matrix governing the large-scale dynamics (Fig. S4).

While Figs. 2-5 examine propagation for an input to V1, which results in propagation along the visual hierarchy, information propagation can be studied along different sensory pathways. In Fig. S5, we test our mechanism on injecting a pulse input to area 2, corresponding to the primary somatosensory cortex, for the connectivity parameter values in Figs. 4B, C. Response activity emerges across a different set of areas, with strong activity in the somatosensory areas in the parietal lobe. Increasing GBA facilitates propagation to the prefrontal areas.

To quantify population synchrony across the regimes investigated in Figs. 4, 5, we examine the global activity in response to a sustained input, based on the asynchronous and synchronous regimes being characterized by stationary and oscillatory population activities respectively [7, 19]. We examine the degree of population synchrony [58] (Fig. S6), based on the variance of the average population voltage of the excitatory population in V1, on moving from the parameter set in Fig. 4B to the parameter set in Fig. 5A (see Methods), and observe a steady increase in the degree of population synchrony.

The stereotypical response onset sequence to short-duration inputs, in the globally active network (Fig. 5B), suggests the existence of an underlying cause. One possibility is that the signal reaches a given area through the shortest possible path. Alternatively, it could reach the area following the path with the strongest connections, even if it is slower than the first case; or could follow a path with strong connections having strengths beyond a threshold. To uncover the actual mechanism, we test these multiple hypotheses (Fig. S7). We observe a close match between the predicted and observed onset times on thresholding the FLNs to include only the strong connections beyond a threshold, and thereby assuming that the signal takes the shortest possible path to reach a given area.

**Fig. S1:**
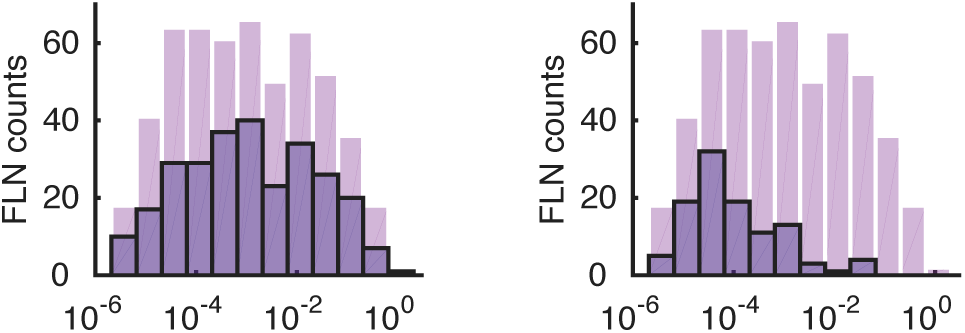
FLN distribution (in light purple) overlaid with feedback FLNs (left) and unidirectional FLNs (right)

**Fig. S2:**
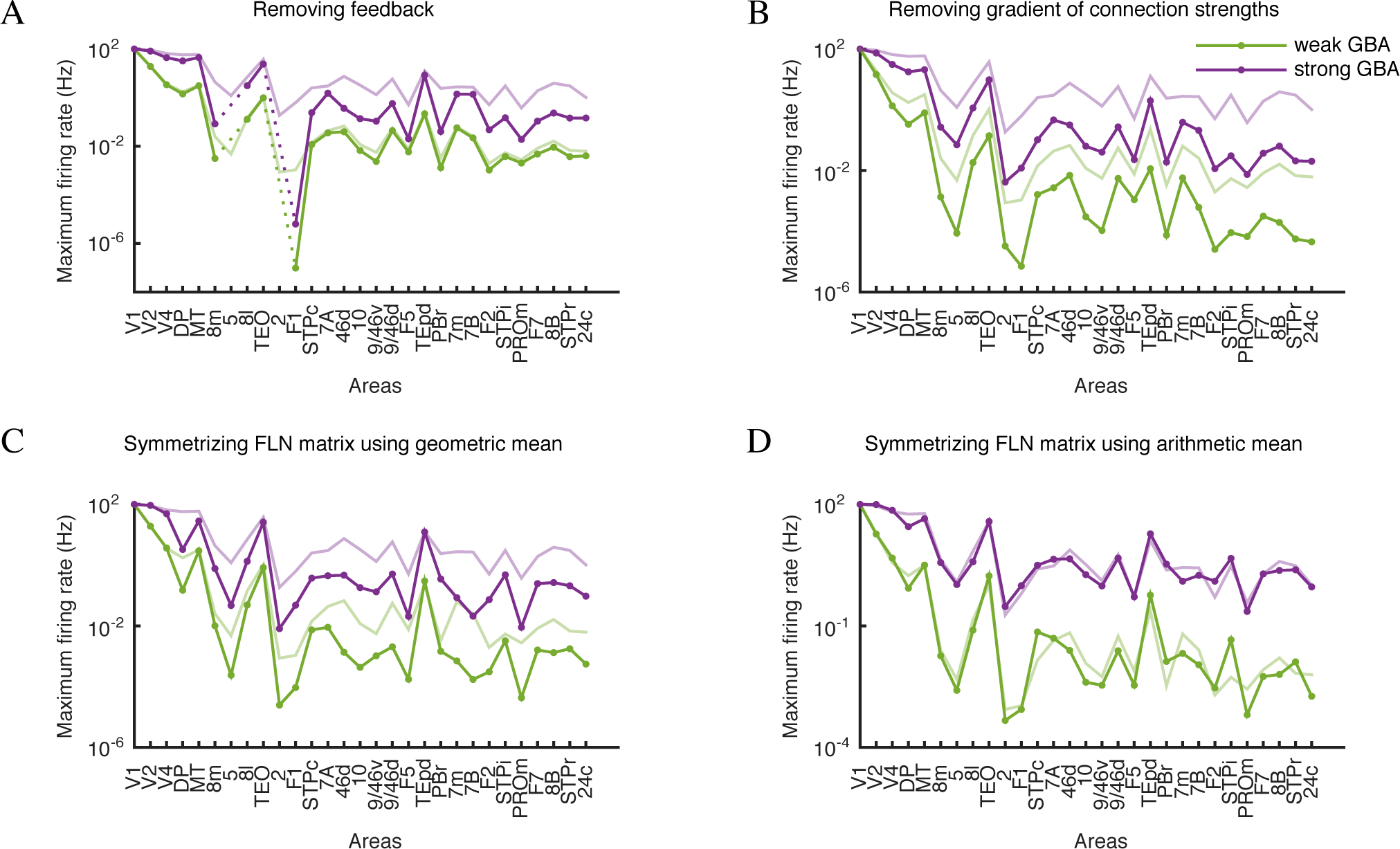
Robustness of the mechanism across different conditions. The light curves show the propagation corresponding to the control case for weak and strong GBA. (A) On removing feedback projections. (B) On removing heterogeneity across areas, that is, setting *η* = 0. (C) On symmetrizing the FLN connectivity matrix using the geometric mean of the inter-areal projection strengths. (D) On symmetrizing the FLN connectivity matrix using arithmetic means.

**Fig. S3:**
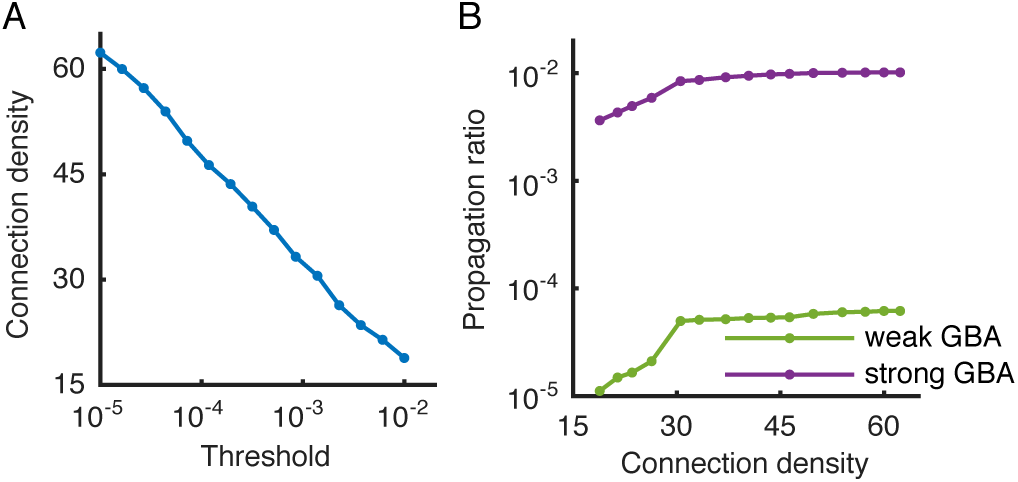
Removing weak anatomical connections. (A) Connection density of the underlying anatomical connectivity matrix as a function of the parametric threshold for which connections below that threshold are removed. (B) Propagation ratio for weak and strong GBA, on removing weak connections.

**Fig. S4:**
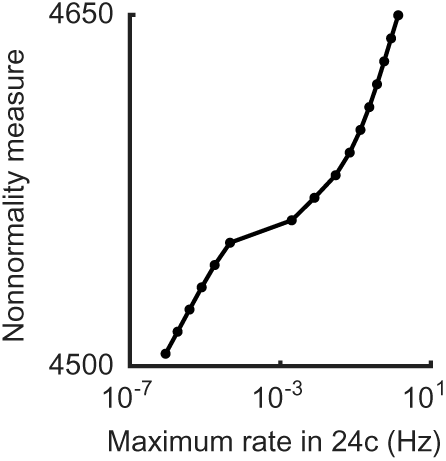
An improvement in propagation is accompanied by an increase in the non-normality measure.

**Fig. S5:**
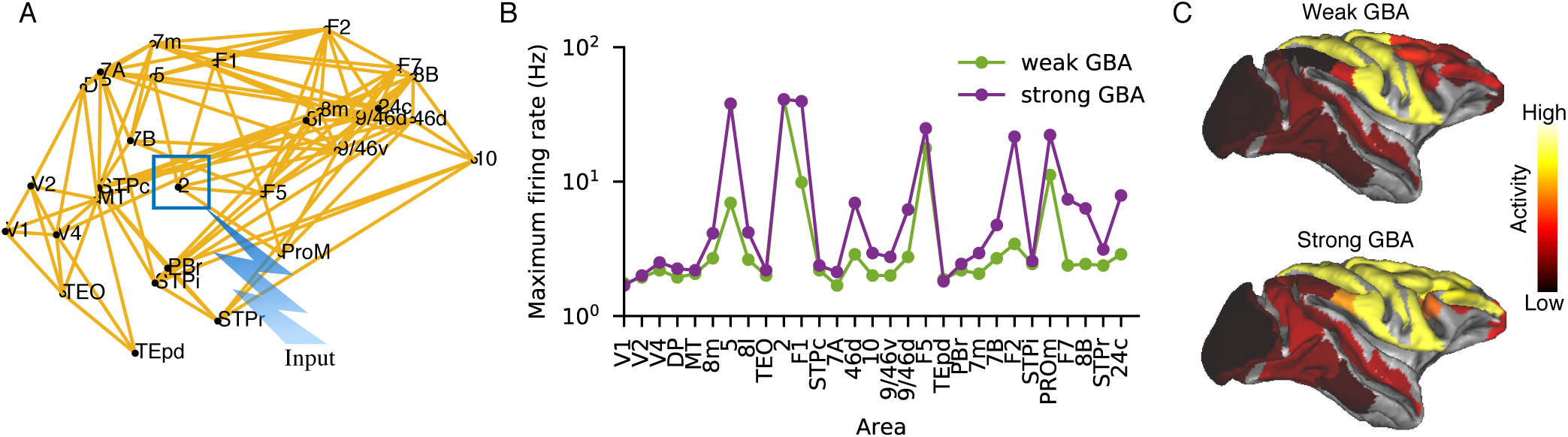
Increasing GBA improves propagation along the somatosensory pathway. (A) Input is injected to the excitatory population in area 2, the primary somatosensory cortex. (B) Peak firing rate across areas as the response to the pulse input to area 2 propagates across the cortical areas for weak (green) and strong (purple) GBA. (C) Lateral plots of the macaque cortical surface for weak and strong GBA, generated using Caret.

**Fig. S6:**
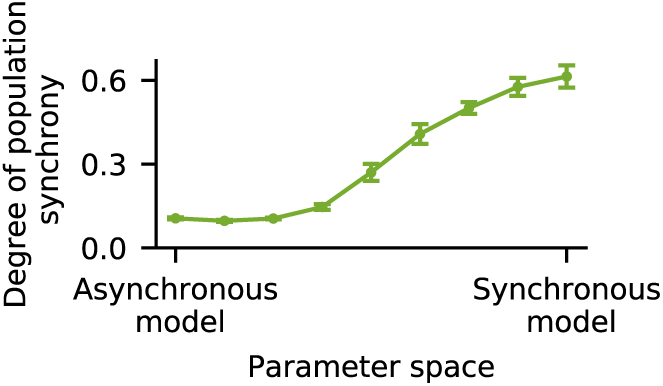
Degree of population synchrony on moving from the parameter set in Fig. 4B to the parameter set in Fig. 5A.

**Fig. S7:**
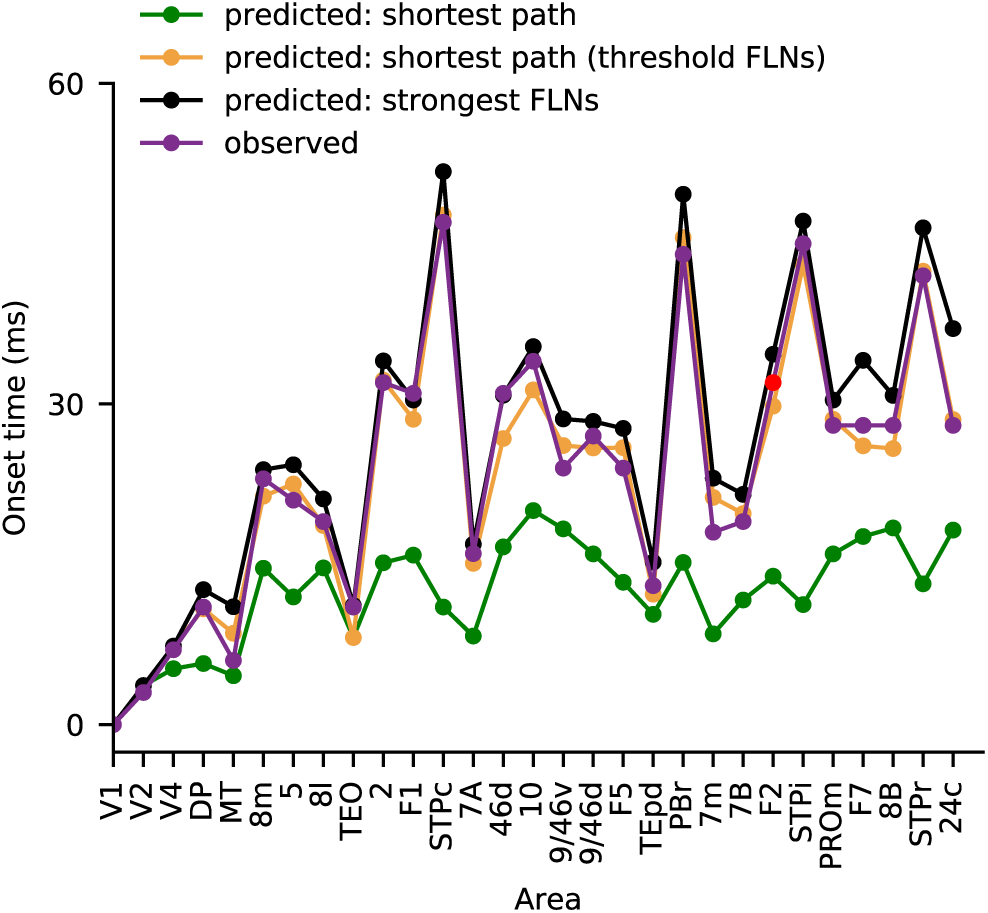
Predicting the observed response onset times across areas as the signal propagates along the hierarchy. The observed onset times are shown in purple. The green curve shows the predicted onset times assuming that the signal starting from V1 follows the shortest possible path to reach a given area. The orange curve is similar to the green, but only considers FLN values with connection strengths exceeding a parametric threshold. The black curve shows the predicted onset times assuming that the signal follows the path over which the sum of FLN values is maximum (that is, the path with the strongest overall connections) to reach a given area.

## Appendix

Consider a local circuit lying in an inhibition-stabilized regime (*w*_*EE*_, *w*_*EI*_ > 1, where *w*_*ij*_ is the connection strength from population type *j* to population type *i*) as in (Fig. 2A), with excitatory and inhibitory firing rates denoted by *E*,*I* respectively. Say the excitatory population is acted upon by a delta-pulse input, to set the initial firing rate to *E*_0_ while *I*_0_ = 0. Without loss of generality, assume *E*_0_ = 1.

The dynamics is described by,

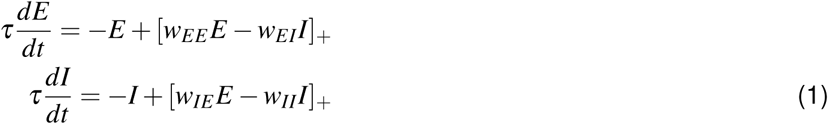

where *x*_+_ = max(*x*,0) denotes rectification.

For fixed *w*_*IE*_, *w*_*II*_, the system stability depends on the local-circuit recurrent excitation wee and the feedback-inhibition *w*_*EI*_. We define the stability boundary as the set of points in the *w*_EE_ – *w*_*EI*_ parameter space for which given the initial condition, the system evolves to a non-zero steady state.

We want to show that,

Given w_*IE*_, *w*_*II*_, the steady-state *E* value on the *w*_*EE*_ – *w*_*EI*_ stability boundary (Fig. 2C) increases on moving along the direction of increasing *w*_*EE*_, *w*_*EI*_.

At steady-state, *dE*/*dt*, *dI*/*dt* = 0.

Say,

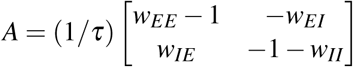

Let *λ*_1_, *λ*_2_ denote the eigenvalues of *A*, and *v*_1_, *v*_2_ be the corresponding eigenvectors. Say,

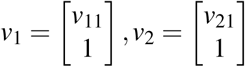

If system 1 were to be linear, then assuming both λ_1_, λ_2_ are not zero,

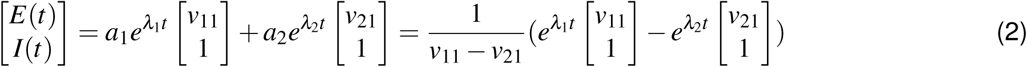

since *E*_0_ = 1, *I*_0_ = 0.

Thus, from 2, whenever *A* has complex conjugate eigenvalues, system 1 is not linear, and the rectification ensures convergence to *E*, *I* = 0. Whenever *A* has a positive real eigenvalue, 1 diverges.

Let *P* be a point in the *w*_*EE*_ – *w*_*EI*_ parameter space corresponding to λ_1_ = 0, λ_2_ < 0. Then, from 2, at *P*,

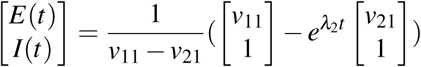

so that system 1 behaves linearly at *P* to converge to

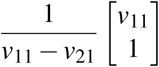

Thus, *P* lies on the stability boundary. (Note that since *w*_*EE*_ > 1, λ_1_, λ_2_ = 0 results in system instability.) Thus, for any point on the stability boundary, 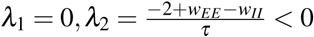.

Now λ_1_ = 0 implies (*w*_*EE*_ – 1)(1 + *w*_*II*_) = *w*_*IE*_*w*_*EI*_, thus, 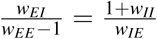. Since we fix *w*_*II*_,*w*_*IE*_, this ratio is constant along the stability boundary.

Say, 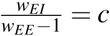 for some constant *c* > 0. Then,

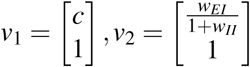

Thus, from 2, the steady-state response at *P* is,

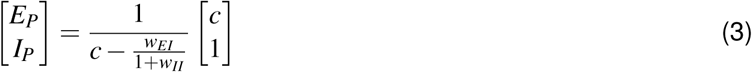

(Note that 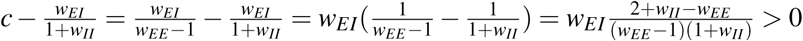 since *w*_*EE*_ > 1, λ_2_ < 0.)

Say *Q* is another point on the stability boundary with parameters 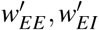 such that 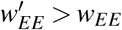 and 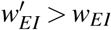. Then, the steady-state excitatory response at *Q*, is 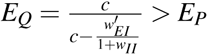.

Thus, moving along the stability boundary in the direction of increasing recurrent excitation (Fig. 2C) shows a progressive increase in the steady-state excitatory firing rate, which can be used to intuitively understand the higher transient amplification prior to decay (Fig. 2B) achievable with stronger recurrent excitation.

